# Small molecule nitroalkenes inhibit RAD51-mediated homologous recombination and amplify triple-negative breast cancer cell killing by DNA-directed therapies

**DOI:** 10.1101/2023.08.11.552990

**Authors:** Lisa Hong, Dennis C. Braden, Yaoning Zhao, John J. Skoko, Fei Chang, Steven R. Woodcock, Crystall Uvalle, Allison Casey, Katherine Wood, Sonia R. Salvatore, Alparslan Asan, Trey Harkness, Adeola Fagunloye, Mortezaali Razzaghi, Adam Straub, Maria Spies, Daniel D. Brown, Adrian V. Lee, Francisco Schopfer, Bruce A. Freeman, Carola A Neumann

## Abstract

Nitro fatty acids (NO_2_-FAs) are endogenously generated lipid signaling mediators from metabolic and inflammatory reactions between conjugated diene fatty acids and nitric oxide or nitrite-derived reactive species. NO_2_-FAs undergo reversible Michael addition with hyperreactive protein cysteine thiolates to induce posttranslational protein modifications that can impact protein function. Herein, we report a novel mechanism of action of natural and non-natural nitroalkenes structurally similar to (*E*) 10-nitro-octadec-9-enoic acid (CP-6), recently de-risked by preclinical Investigational New Drug-enabling studies and Phase 1 and Phase 2 clinical trials and found to induce DNA damage in a TNBC xenograft by inhibiting homologous-recombination (HR)-mediated repair of DNA double-strand breaks (DSB). CP-6 specifically targets Cys319, essential in RAD51-controlled HR-mediated DNA DSB repair in cells. A nitroalkene library screen identified two structurally different nitroalkenes, a non-natural fatty acid [(*E*) 8-nitro- nonadec-7-enoic acid (CP-8)] and a dicarboxylate ester [dimethyl (*E*)nitro-oct-4-enedioate (CP- 23)] superior to CP-6 in TNBC cells killing, synergism with three different inhibitors of the poly ADP-ribose polymerase (PARP) and γ-IR. CP-8 and CP-23 effectively inhibited γ-IR-induced RAD51 foci formation and HR in a GFP-reported assay but did not affect benign human epithelial cells or cell cycle phases. In vivo, CP-8 and CP-23’s efficacies diverged as only CP-8 showed promising anticancer activities alone and combined with the PARP inhibitor talazoparib in an HR-proficient TNBC mouse model. As preliminary preclinical toxicology analysis also suggests CP-8 as safe, our data endorse CP-8 as a novel anticancer molecule for treating cancers sensitive to homologous recombination-mediated DNA repair inhibitors.

**Graphical Abstract:** 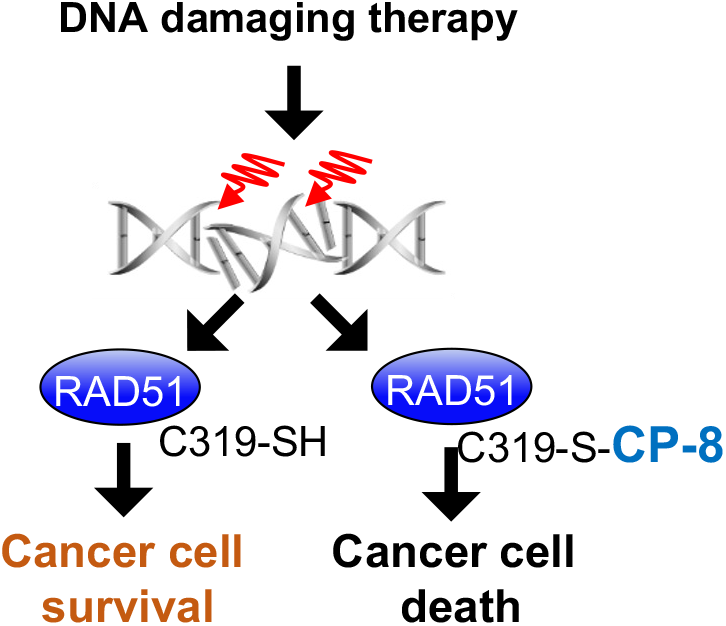

## INTRODUCTION

Covalent inhibitors have enjoyed a long history in drug discovery that began with aspirin in the late 19^th^ century, was followed by penicillin and omeprazole in the twentieth century, and is now culminating with a recent surge of rationally engineered kinase inhibitors. Covalent inhibition entails a bond-forming event of a small molecule with a protein that can be reversible or irreversible. While many acrylamides (e.g., ibrutinib, osimertinib, afatinib) and epoxide-containing drugs (e.g., carfilzomib, fosfomycin) form irreversible bonds with their protein targets, electrophilic drugs, such as electron-deficient ketones, enoates, nitriles, or nitroalkenes, form reversible covalent bonds with their respective targets. This reversibility stems from an additional electron withdrawing group at the alpha carbon of the Michael acceptor^1^.

Nitroalkene fatty acid (NO_2_-FA) derivatives are generated endogenously from unsaturated fatty acids in the presence of reactive oxygen and nitrate species. NO_2_-FAs can be detected in several tissues and biofluids and signal as adaptive metabolic and inflammatory mediators and inhibitors of pathogenic cell proliferation^2^. NO_2_-FAs with a nitroalkene substituent are reactive electrophiles that facilitate reversible adduction to a small population of highly reactive cysteine thiolates via Michael addition ^3^. At physiological and therapeutic concentrations, nitro-oleic acid (OA-NO_2_) predominantly alkylates intracellular glutathione (GSH) and susceptible protein cysteine residues, thereby impaction protein structure and enzyme catalytic function ^3^. For example, OA-NO_2_ (CP-6) canonical protein cysteine targets include Cys285 in the nuclear lipid receptor peroxisome proliferator-activated receptor γ (PPARγ), Cys38 in NF-κB, and Cys273 and 288 in the Kelch-like ECH-associated protein 1 (Keap1) regulator of nuclear factor (erythroid-derived-2)-like2 (NRF2) ^4–6^.

We recently demonstrated that CP-6 inhibits viability, migration, and invasion of triple-negative breast cancer (TNBC) cell lines in vitro and decreases tumor growth of a human TNBC xenograft in mice ^7^. In this study, we provided evidence that CP-6 impeded NF-κB signaling in several ways through (*a*) blocking of the inhibitor of NF-κB subunit kinase β phosphorylation and downstream inhibitor of NF-κB degradation, (*b*) adducting the NF-κB RelA protein to prevent DNA binding, and (*c*) by promoting RelA polyubiquitination and proteasomal degradation^7^. Exploring other CP-6 effects on cell viability, we noticed it strongly induces DNA damage in the form of double-strand breaks (DSB) in TNBC cell lines in vitro and tumor tissues of a human TNBC xenograft model, suggesting that CP-6 may directly perturb the DNA damage response. In parallel, we discovered that the recombinase RAD51, known to function in concert with auxiliary mediator proteins to direct the repair of DNA DSBs via homologous recombination (HR) and a sought-after drug target in cancer therapy ^8, 9^, contains a redox-sensitive cysteine (Cys319) with nucleophilic characteristics. Cys319 is essential for RAD51 functions in cells, and the oxidation of the Cys319 thiol negatively impacts RAD51 binding to single-strand (ss) DNA ^10^, a prerequisite for RAD51-mediated repair of DNA DSB. As others have shown that Cys319 in RAD51 is solvent-exposed and available for alkylation ^11^, we investigated if CP-6 adducts to Cys319 and affects RAD51 function. We found a specific reaction of CP-6 with RAD51 Cys319 and further identified that combination treatment of CP-6 with anti-neoplastic agents such as doxorubicin, cisplatin, IR, or olaparib enhances the antiproliferative effect of these DNA-damaging agents in TNBC cells.

Consequently, in TNBC cells, CP-6 suppressed IR-induced RAD51 foci formation, HR, RAD51 binding to ssDNA, RAD51-ABL interaction, and RAD51 Tyr315 phosphorylation, as well as induced phosphorylation of Ser139 H2AX (lJH2AX) ^7^. Our observations reinforced the concept that NO_2_-FAs such as CP-6 disrupt HR and reveal a novel therapeutic strategy where redox- derived soft electrophile species can, in turn, sensitize cancer cells to DNA-directed therapeutic strategies. There is a broad interest in developing small molecule RAD51 inhibitors. First-generation RAD51 inhibitors such as B02, RI-1, RI-2, and IBR2 are limited by a poor potency for growth inhibition, toxicity at the high micromolar inhibition concentrations (IC50) required in cell test systems that is also in part due to irreversible reactions with RAD51 and bystander proteins^12^.

Poly-ADP ribose polymerase (PARP) inhibitors are FDA-approved for treating TNBC mutants for *BRCA1* or *BRCA2* genes. As BRCA1/2 genes are essential in repairing DNA DSBs via homologous recombination, loss of function mutations renders TNBC cells sensitive to DNA damaging agents such as PARPi. However, genes ∼80% of all TNBC patients express wildtype BRCA1/2 genes and are therefore not sensitive to PARPi treatments. Therefore, as we previously showed that CP-6 inhibits RAD51 function and sensitizes BRCA1/2 wildtype TNBC cell lines to IR and PARPi treatment, we compared CP-8 and CP-23 for synergism with different PARPis that vary in potency. Some evidence suggests the potency of a PARPi is correlated with its ability to trap PARP at the site of the lesion ^13, 14^. For example, talazoparib’s trapping potency is ∼10,000-fold greater than other PARPi, while its catalytic potency only differs ∼ 40-fold ^15^. Therefore, we decided to evaluate the synergy of our NFA candidates with multiple PARPi with varying PARP-trapping abilities. PARPi DNA trapping potency can be ranked as follows: *talazoparib**>>**niraparib>olarapib/rucaparib* ^16^.

To optimize RAD51 inhibition by small molecule nitroalkenes, a library of 60 candidate leads was screened for RAD51 inhibition and tumor cell killing. Of this library, two lead compounds CP-8, [(*E*) 8-nitro-nonadec-7-enoic acid] and CP-23, a dicarboxylate ester [dimethyl (*E*)nitro-oct- 4-enedioate] were chosen for further testing. There was a striking impact of CP-8, with TNBC cell killing and tumor volume reduction, both as a single agent and combined with the poly-ADP ribose polymerase (PARP) inhibitor talazoparib. At the same time, CP-23 showed no efficacy in vivo. The pharmacokinetic analysis detected significant levels of inactive CP-23 metabolites in murine plasma and mammary tumor tissue, partly explaining the lack of in vivo efficacy. In contrast, CP-8 was detected predominantly in its unmetabolized form in plasma, benign and tumorous mammary glands, and brain tissue and showed no toxicity to the bone marrow, encouraging the further preclinical evaluation of CP-8 as a reversible non-toxic inhibitor of RAD51 in HR-mediated DNA DSB repair.

## RESULTS

The rationale for designing and synthesizing the nitroalkene structural variants tested herein was a) initial molecular dynamic modeling suggested that closer proximity of the nitroalkene to the carboxylate-terminus would enhance nitroalkene (NA) association with the RAD51 C-terminal groove, b) CP-1 through CP-22 and CP-23 through CP-31 were homologs of endogenous nitroalkene metabolites with or without hydrophobicity-conferring esterified carboxylates. **Figure 1** shows that CP-8, compared with OA-NO_2_ (CP-6), has the nitroalkene placed two carbons closer to the carboxylate terminus and one carbon longer than CP-6.

**Figure 1.**
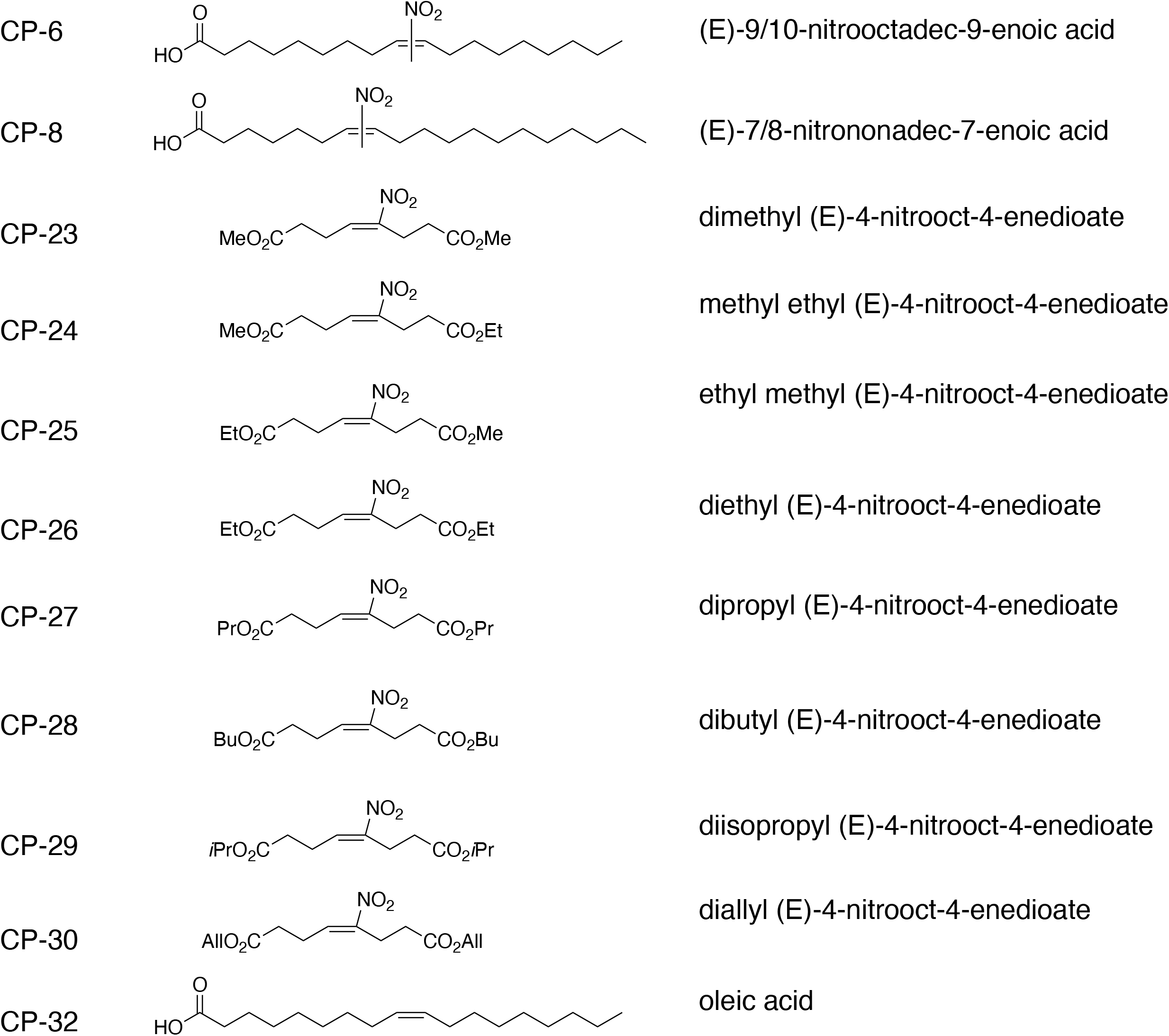
Chemical structures of example compounds.

### Nitroalkenes inhibit TNBC cell growth

Growth inhibitory effects of 60 compounds were evaluated in human and murine TNBC and benign human breast epithelial cells (**Figure 2**). We have previously reported that CP-6 inhibited the growth of TNBC cell lines expressing wildtype genes of *BRCA1* and *BRCA2* (MDA-MB-231, BT-549, and Hs578t) ^7,17^. To expand the screening analysis, we included two human immortalized breast epithelial cell lines (MCF-10A, hTERT-HME1) and one murine TNBC cell line (4T1) that expresses wildtype *BRCA1* and *BRCA2* genes ^18^. Cells were treated with nitroalkenes on days 0, 2, and 6. On day 7, growth inhibition was quantified by measuring the ATP-dependent luminescent signal generated using Ultra-Glo luciferase with the substrate luciferin. Strikingly, CP-27 to CP-30 displayed minimal growth inhibitory effects on MDA-MB-231 cells, with EC50 values substantially higher than 10 μM. CP-8, on the other hand, displayed the lowest EC50 value of 1.79 μM (±0.17 μM), while CP-6 exhibited a value of 3.96 μM (±0.16 μM), CP-24 with 3.4 μM (±0.24 μM), CP-23 with 3.6 μM (±0.14 μM), CP-25 with 4.45 μM (±0.27 μM) and CP-26 with 5.22 μM (±0.32 μM) (**Figure 2A**). Interestingly, NA growth inhibitory effects on benign cells varied, where CP-6 generated an EC50 value of 2.15 μM (±0.01 μM) in hTERT- HME1 cells but an EC50 value substantially higher than 10 μM in MCF-10A cells. Similarly, CP- 23, CP-24, and CP-25 showed lower EC50 values in hTERT-HME1 cells with values of 9.13 μM (±1.98 μM), 4.67 μM (±0.10 μM), and 5.42 μM, respectively (**Figures 2B and 2C**). We compared CP-8 in another TNBC cell line, MDA-MB-468, that, like MDA-MB-231 cells, expresses wildtype *BRCA1* and *BRCA2* genes^17^. CP-8 EC50 values in both cell lines were comparable at 1.79 (±0.19 μM) for MDA-MB-231 and 1.71 (±0.16 μM) for MDA-MB-468 cells (**Figure 2D**). Notably, the difference in the EC50 values of both TNBC cell lines, when compared to EC50 values of MCF-10 cells, was highly significant (p<0.0001). Notably, CP-8 EC50 values increased ∼100-fold from 4.9 μM ±0.32 μM) to 489.5 μM (±1377 μM) when cells were plated at 10-fold higher density (**Figure 2D**). Lastly, comparing CP-8 and CP-23 EC50 values in the murine TNBC cell line, 4T1 showed a slightly lower EC50 value for CP-23 with 2.19 μM (±0.27 μM) compared to CP-8 with 3.37 μM (±0.51 μM) (**Figure 2E**).

**Figure 2.**
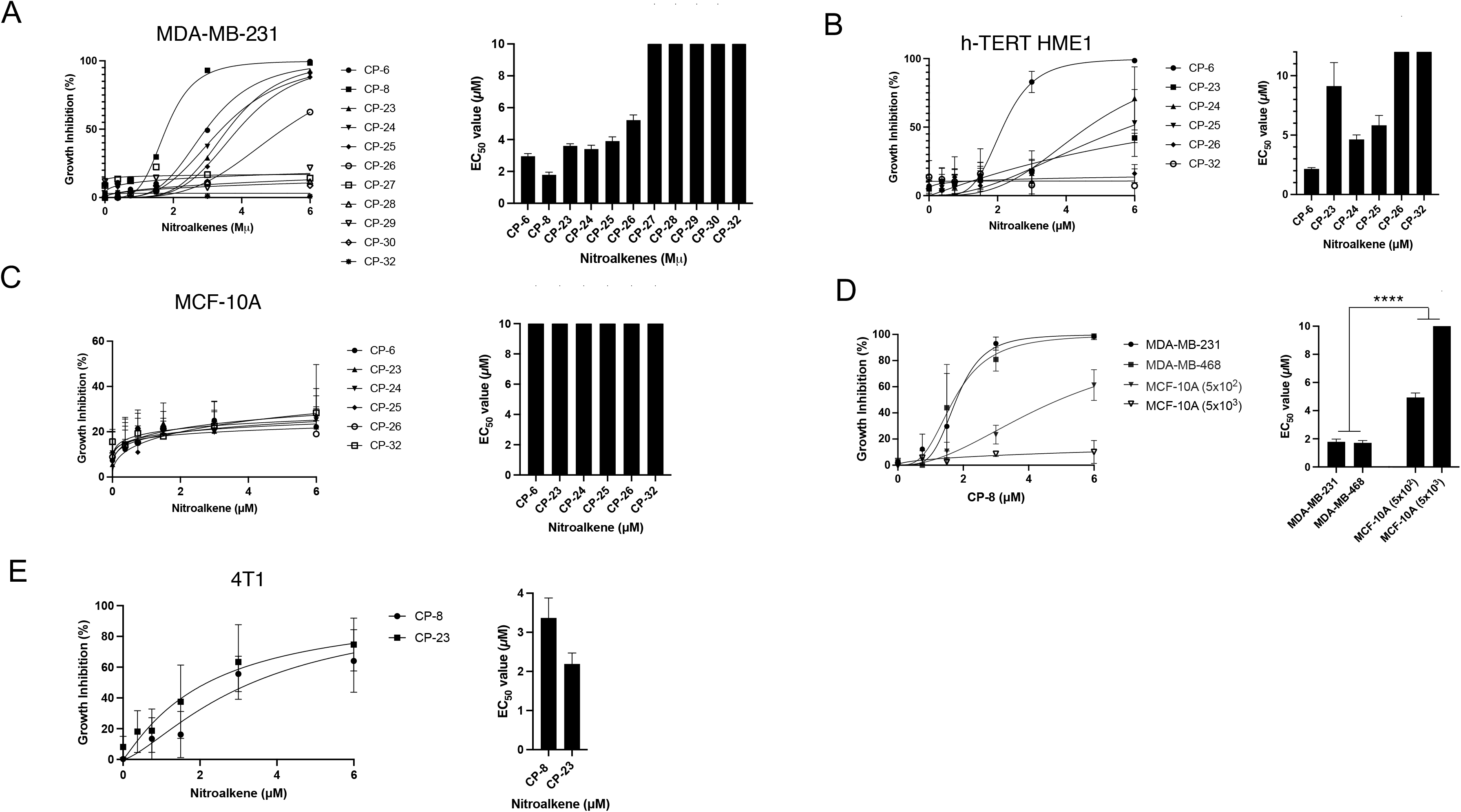
The nitroalkene analogs CP-8 and CP-23 exhibit differential growth inhibition of cancer and normal cells. **A**. MDA-MB-231, **B.** hTERT HME1, **C.** MCF10A or **D.** MDA-MB-231, MDA-MB-468, MCF-10 plated subconfluently (5x10^2^) and confluently (5x10^3^), **E.**4T1 cells were plated in a 96-well plate (500 cells/well) and dosed with 0 to 6 µM nitroalkene analogs on days 0, 2 and 6. Growth inhibition was determined by measuring relative ATP levels on day 7 via CellTiter-Glo (Average ± SEM n=3 except for CP-27 to CP-30 (n=1)).

### CP-8 and CP-23 act synergistically with PARP inhibitors

Both CP-8 and CP-23 enhanced PARPi-induced growth inhibition and acted synergistically with PARPis. In MDA-MB-231, we observed the CP-8 EC50 value (1.79 μM ±0.17 μM) to be ∼60% lower than the CP-23 EC50 value (4.37 μM ±0.19 μM) (**Figure 3A**). CP-8 was more efficacious in enhancing niraparib and olaparib-induced growth inhibition compared to CP-23: addition of CP-8 to olaparib’s EC 50 (2.87 μM ±0.31 μM) or niraparib’s EC50 (2.80μM ±0.37 μM) increased growth inhibition in MDA-MB-231 cells at lower concentrations compared to CP-23, whereas the effect of CP-8 on talazoparib growth inhibition was only slightly more than the one observed after CP-23 addition (**Figures 3B-3D and Supplemental Figures 1A-1F**). Comparing the potency of CP-8 and CP-23 in lowering PARPi EC50 values showed that less CP-8 was needed to lower the EC50 values of all three PARPi significantly. For example, CP-8 significantly lowered olaparib’s EC50 value at 1.5 μM, while 6 μM of CP-23 was needed to reduce the olaparib EC50 value (**Figures 3E-3F**) significantly. Relative CP-8 and CP-23 effects on talazoparib’s EC50 were similar: 1.5 μM of CP-8, compared to 3.0 μM of CP-23, were needed to obtain significant talazoparib EC50 value reduction (**Figures 3E-3F**). Notably, the highest single agent model (HSA; synergyfinder.fimm.fi) predicts the synergy of CP-8 and CP-23 with all three PARPis (**Table 1**). CP-8 addition in the 0.8-3 μM range was synergistic with olaparib (0.8- 12.5μM), niraparib 0.2-3.1 μM, and talazoparib (0.01 to 0.2 μM), suggesting that dose reduction of all three PARPis is possible through CP-8 cotreatment. Importantly, CP-8 exhibited the highest synergy with niraparip and talazoparib at concentrations ∼10-fold lower than niraparib’s and talazoparib’s EC50 values, and ∼8-fold for olaparib’s EC50 values, respectively (**Figure 3G**). Synergies with CP-23 suggested a possible dose reduction for olaparib when CP-23 was added between 3-6 μM. More clear results were obtained for CP-23 cotreatment and niraparib, where 0.8 to 6μM synergized with low niraparib dosing (0.8 to 12.5μM) and even lower for talazoparib (0.01 to 0.2μM). Like CP-8, CP-23 exhibited the highest synergy with talazoparib at concentrations ∼10-fold lower than talazoparib’s EC50 values. In contrast to CP-8, CP-23’s synergy with niraparib lowered its EC50 only about 3.5 fold (**Figure 3H**).

**Figure 3.**
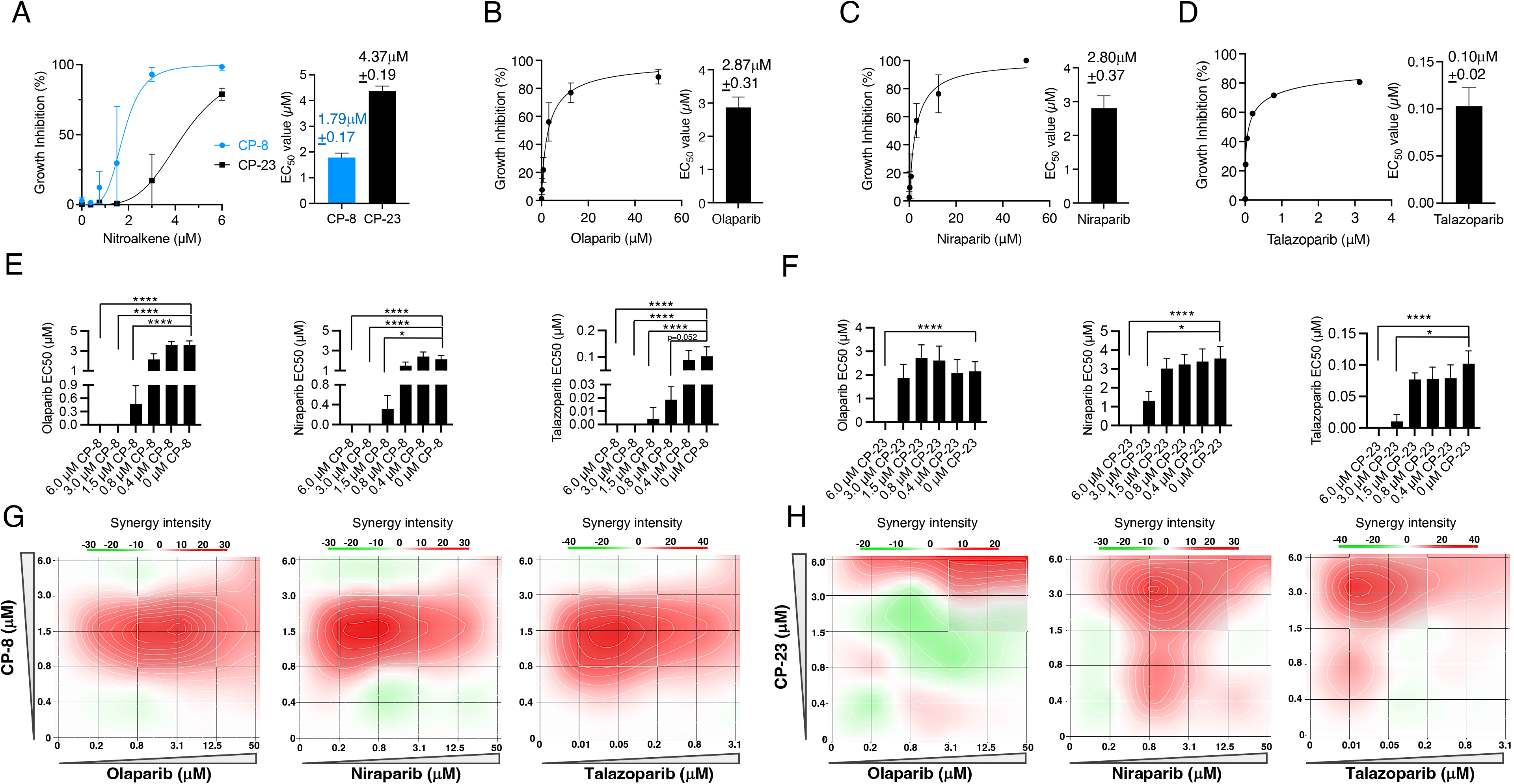
CP-8 and CP-23 display synergistic antiproliferative activity in combination with olaparib, niraparib and talazoparib. A. MDA-MB-231 cells (500 cells/well) were treated with nitroalkene analogs (0 to 6 µM) on days 0, 2, and 6. Growth inhibition was determined by measuring relative ATP levels on day 7 via CellTiter-Glo. **B-D.** MDA-MB-231 cells (500 cells/well) were treated with PAPR inhibitors olaparib, niraparib, and talazoparib on days 0, 2, and 6. EC50 values were determined as in (A). **E-F.** Decrease of EC50 values of PARPi co- treated with CP-8 or CP-23 in increasing concentrations. Statistical analysis was done by two- way ANOVA; Average ± SEM n=3. **G-F.** Drug combination synergy calculations were performed using the SynergyFinder and HSA methods. MDA-MB-231 cells plated as in A-D were treated with 0-6 μM of nitroalkene and increasing doses of PARP inhibitors (olaparib 0-50 μM, niraparib 0-50 μM, and talazoparib 0-3.1 μM). Average ± SEM n=3.

**Table 1:** Synergy scores for CP-8 and CP-23 with individual PARPi generated with the HSA method.

| Drug Combination | Synergy Score | Most Synergistic Area Score | Method |
| --- | --- | --- | --- |
| Olaparib + CP-8 | 8.44 | 16.59 | HSA |
| Niraparib + CP-8 | 5.36 | 10.05 | HSA |
| Talazoparib + CP-8 | 4.39 | 11.41 | HSA |
| Olaparib + CP-23 | 5.23 | 9.98 | HSA |
| Niraparib + CP-23 | 1.3 | 5.28 | HSA |
| Talazoparib + CP-23 | 5.06 | 9.44 | HSA |

### CP-8 and CP-23 differentially sensitize only MDA-MB-231 cells but not MCF-10A cells to IR

Given previous evidence that nitroalkenes selectively sensitize TNBC cells to ionizing radiation (IR)^6^ and the importance of IR as frontline therapy in the clinic, we wanted to test the relative radio-sensitizing capabilities of our two lead candidates, CP-23 and CP-8, by supplementing MBA-MB-231 cells and MCF10A benign breast cells with our candidates and evaluating colony formation ability 7 days after exposure to various doses of IR. Surprisingly, while CP-8 significantly radio-sensitized MBA-MB-231 cells to IR as expected (∼60% reduction in apparent IR IC_50_ value) (**Figure 4A**), CP-23 only very weakly sensitized MBA-MB-231 cells to IR (∼26% reduction in apparent IR IC_50_ value) and the magnitude of this sensitization approached, but did not reach, statistical significance (**Figure 4C**). CP-8 and CP-23 supplementation had similarly disparate effects in MCF10A cells. Namely, CP-8 significantly protected MCF10A cells from IR (∼41% increase in apparent IR IC50 value), while CP-23 did not alter the radiosensitivity of MCF10A cells (**Figure 4B and 4D**). This suggests that CP-8 exhibits better selective radio sensitization properties than CP-23.

**Figure 4:**
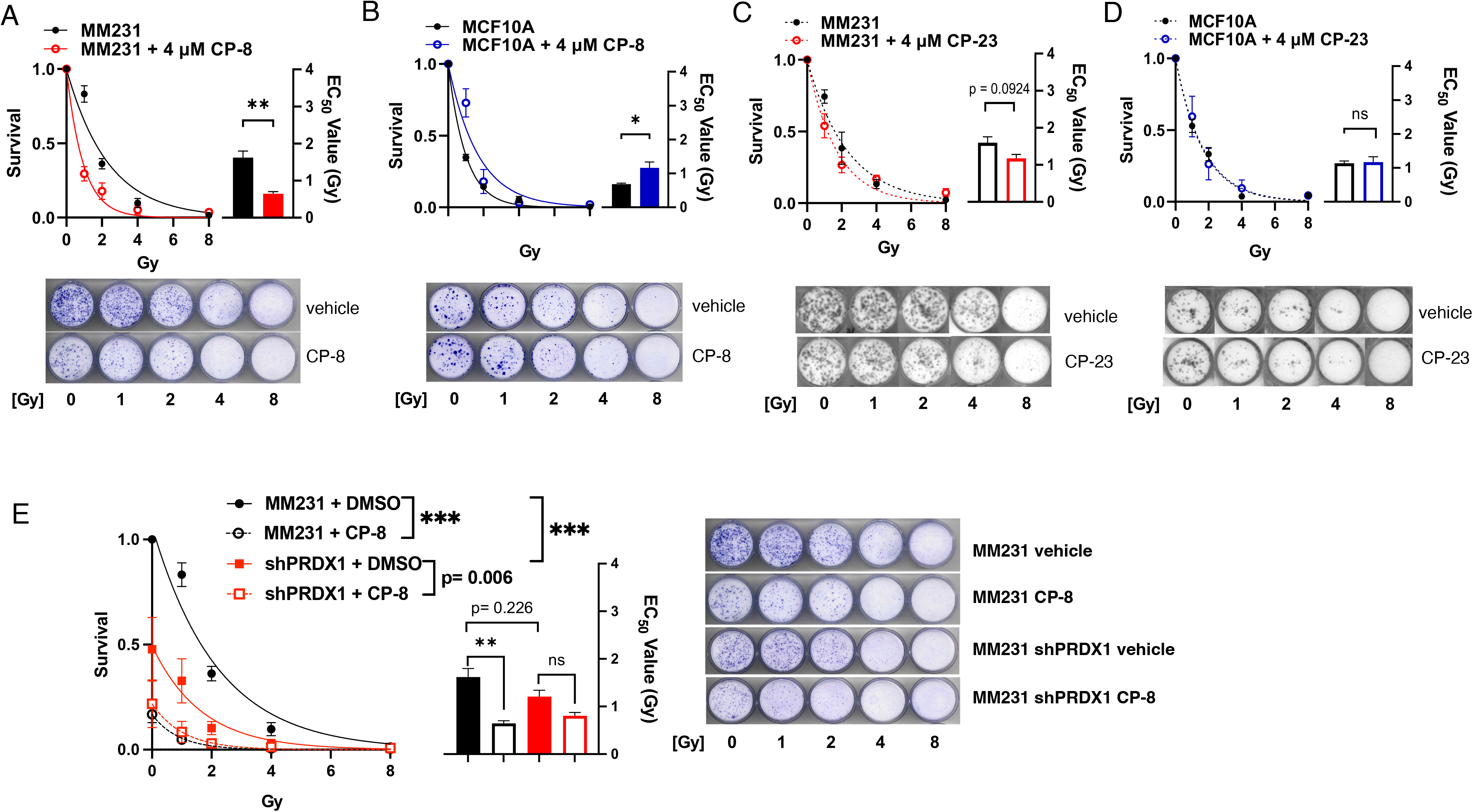
CP-8 has superior cysteine oxidation-dependent radio-sensitizing properties to CP-23. MDA-MB-231 and MCF10A cells were seeded at low density, and subjected to various doses of IR in the presence of either DMSO, CP-23, or CP-8, and colony formation was analyzed after 7 days by crystal violet staining. Colony formation was quantified as survival relative to respective 0Gy controls after supplementation with either CP-8 (**A-B**) or CP-23 (**C-D**). Data plotted as Average ± SEM (N=3) and fit to a one-phase exponential decay function to calculate EC50 values. Statistical significance was determined by a two-tailed student’s t-test. **E.** PRDX1 WT or shPRDX1 MBA-MB-231 cells were irradiated as in (A-D) in the presence of either DMSO or CP-8. Colony formation was quantified as survival relative to PRDX1 WT 0Gy control. Data plotted as Average ± SEM (N=3) and fit to a one-phase exponential decay function. Statistical significance was determined by two-way ANOVA followed by Tukey’s test for multiple comparisons of AUCs. Representative images for all colony formation experiments are included in each panel. ***, p < 0.001, **, p < 0.01, *, p < 0.05.

We recently demonstrated that loss of the redox regulatory protein Peroxiredoxin 1 (PRDX1) oxidizes RAD51 Cys319 and inhibits DSB repair analogous to nitroalkenes^9^. Therefore, we wanted to know whether constitutively mimicking RAD51 Cys319 oxidation by knocking down PRDX1 would be epistatic with treatment with nitroalkenes on IR sensitization. To test this hypothesis, we supplemented PRDX1 WT or PRDX1 shRNA knockdown MBA-MB-231 cells with CP-8 and evaluated colony formation ability as described previously. As expected, the knockdown of PRDX1 alone significantly sensitized MBA-MB-231 cells to IR (∼61% reduction in AUC) compared to untreated PRDX1 WT cells. Further supplementation with CP-8 only marginally increased the radiosensitivity of MDA-MB-231 cells (∼87% reduction in AUC) compared to untreated PRDX1 WT cells, and the magnitude of this increase did not reach statistical significance (**Figure 4E**). Therefore, we conclude that PRDX1 knockdown is at least partially epistatic with CP-8 treatment. This finding further supports the hypothesis that the mechanism of nitroalkene toxicity is partially mediated by RAD51 Cys319 alkylation.

### CP-8 and CP-23 inhibit HR efficiency and reduce RAD51 foci formation

Cell cycle analysis of MDA-MB-231 cells confirmed that while CP-23 did not induce significant changes to the cell cycle, CP-8 slightly increased the number of cells in G2 and decreased the number of cells in the G1 phase of the cell cycle (**Figures 5 A and B**). CP-8 and C-23- dependent effects on HR DNA repair were compared using a DR-GFP reporter assay and immunofluorescence staining of RAD51 foci, a clinically recognized biomarker for HR efficiency ^19–23^. This analysis quantifies the intracellular recombination of an integrated cDNA cassette of two tandem non-fluorescent GFP constructs following the introduction of an I-SceI cleavage to the system by measuring the fluorescent GFP protein produced following successful recombination ^24^. Daily treatment of U2OS cells with either CP-8 or CP-23 harboring the DR-GFP construct revealed that after I-SceI transfection, the number of GFP positive cells was significantly decreased by ∼ 37% with 1μM CP-8 compared to native OA after 48 h (**Figure 5C**). In contrast, 1μM CP-23 had minimal effect on HR, and 5μM reduced HR by ∼46% (**Figure 5D**). Notably, both CP-8 and CP-23 (1μM) inhibited 5Gy-induced RAD51 foci formation in MDA-MB- 231 cells similarly by ∼65% and ∼68%, respectively (**Figures 5E-5F**).

**Figure 5.**
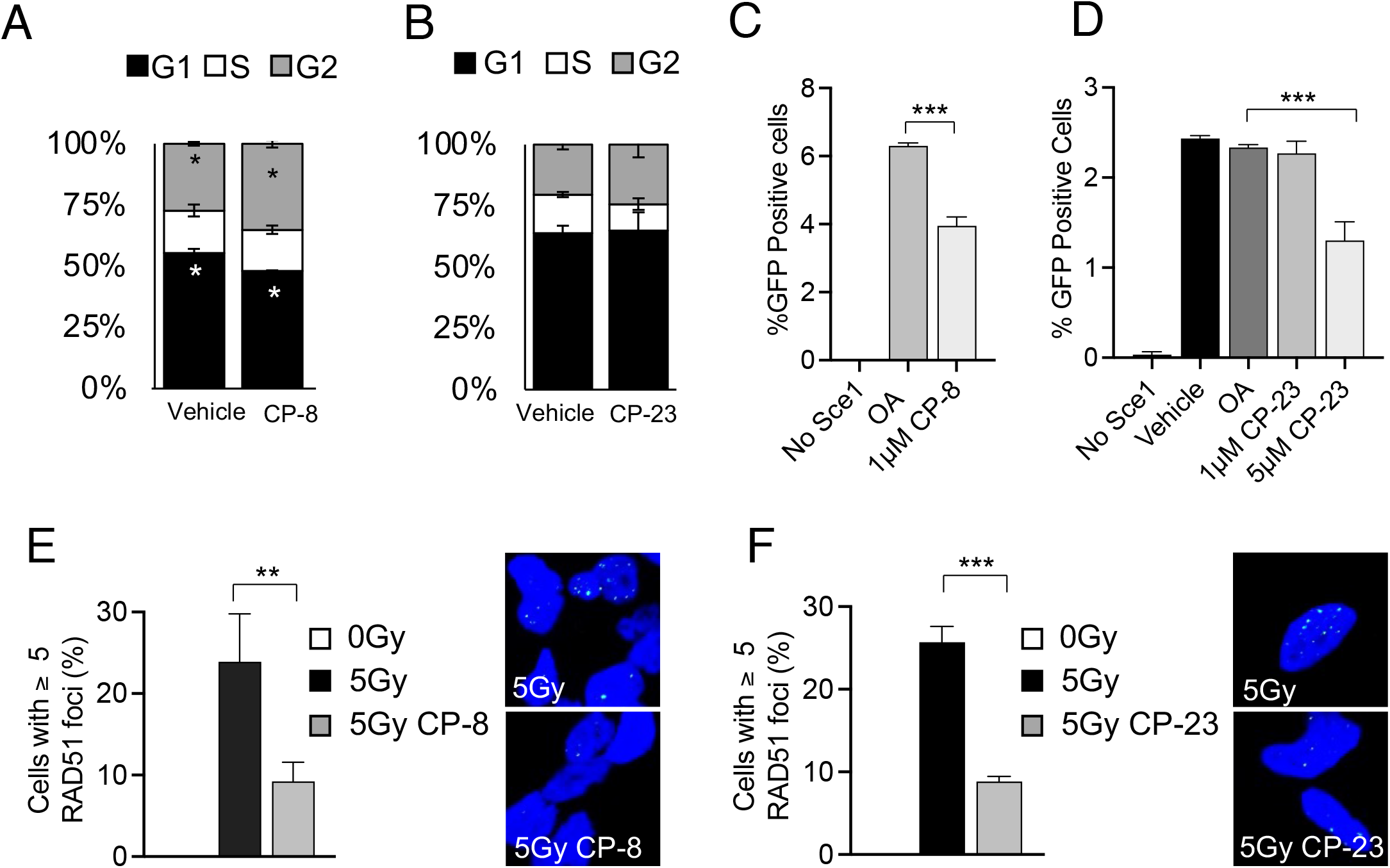
CP-8 inhibits HR more potently than CP-23. A and B. MM231 cells were treated with vehicle or 4 µM CP-8 or CP-23 for 24 h and then processed for cell cycle analysis (propidium iodide DNA staining) with flow cytometry. Mean ± SD, N=3. **C and D.** U2OS cells containing the HR reporter construct DR-GFP were transfected with an I-SceI plasmid and treated with vehicle (gray) 1.3 μM of OA, CP-6, CP-8, or CP-23. Negative control cells did not have I-SceI present. Vehicle (DMSO) and oleic acid (OA) were included as controls. Average + SEM, n ≥ 3. **E and F**: 1.3 µM CP-8 or CP-23 treatment effects on RAD51 nucleoprotein formation after 5 Gy ionizing radiation exposure. MDA-MB-231 cells (150,000 cells/coverslip) nitroalkene 30 min after dosing cells with 5 Gy inhibited irradiation-induced RAD51 foci formation as detected by immunofluorescent confocal microscopy. Cells were processed 6 h after IR, images were acquired using a Nikon A1R confocal microscope with 60× oil objective, and the acquisition was performed using NIS Elements software. Quantification of z-stacks and foci was completed using ImageJ software—average + SEM n=3. One-way ANOVA was used to test the significance.

### CP-8 but not CP-23 inhibits *in vivo* TNBC tumor growth

To compare the anti-tumor effect of both nitroalkenes in vivo, a commonly used human TNBC xenograft model was used where MDA-MB-231 cells were transplanted into the mammary gland of nude mice. Mice were dosed daily when tumors reached about 40 mm^3^ in size and with CP-8 (**Figure 6**) or CP-23 (**Supplemental Figure 2**) and PARPi (talazoparib and olaparib, respectively). CP-8 (15mg/kg, PO) decreased TNBC tumor burden as a single agent and in combination with low-dose PARPi tala (0.1 mg/kg, PO compared to 0.3mg/kg in other studies ^25–28^) (**Figure. 6A**). In contrast, neither 15 mg/kg nor 50 mg/kg, CP-23 PO decreased TNBC tumor burden as a single agent treatment or offered dramatic tumor diminishment in combination with olaparib (50 mg/kg, PO) (**Supplemental Figure 2A**). Thus, we stopped this experiment after 21 days. Only CP-8 alone or combined with talazoparib showed tumor volume reduction (**Figure 6B**). We observed the complete disappearance of three established tumors in each arm (**Figure 6C**). Mouse weights were not affected by either treatment (**Figure 6D**). To better understand why CP-23 did not affect tumor reduction, a pharmacokinetic (PK) analysis was performed for 1 wk. Mice were dosed once with 100 mg/kg CP-23 PO and by mass spectrometry. This analysis revealed that most CP-23 is metabolized in vivo by oxidation reactions leading to di- carboxylated and mono-methyl/mono-carboxylated non-electrophilic CP-23 metabolites (**Supplemental Figure 2B**). In contrast, after one dose of CP-8, there was 0.01 to 0.02 pmol/mg tissue native CP-8 detectable in the mammary gland, mammary tumor, and brain tissue. Increasing CP-8 dosing to 50 mg/kg increased tissue concentrations up to 0.04 pmol/mg tissue, while plasma levels remained in the 2.3 nM to 5.2 nM range (**Figure 6E**). Fatty acid β- oxidation metabolites of CP-8 (dinor and tetranor) and alkene-reduced forms of CP-8 and related dinors and tetranors, both esterified and free, were identified in tissues and plasma. These concentrations increased ∼3-fold when the 15 mg/kg dose was increased to 50 mg/kg (**Supplemental Figures 4A-4G**). Notably, the weight of 15 mg/kg and 50 mg/kg treated cohorts did not change compared to vehicle controls for 21 days (**Supplemental Figure 4G**).

**Figure. 6.**
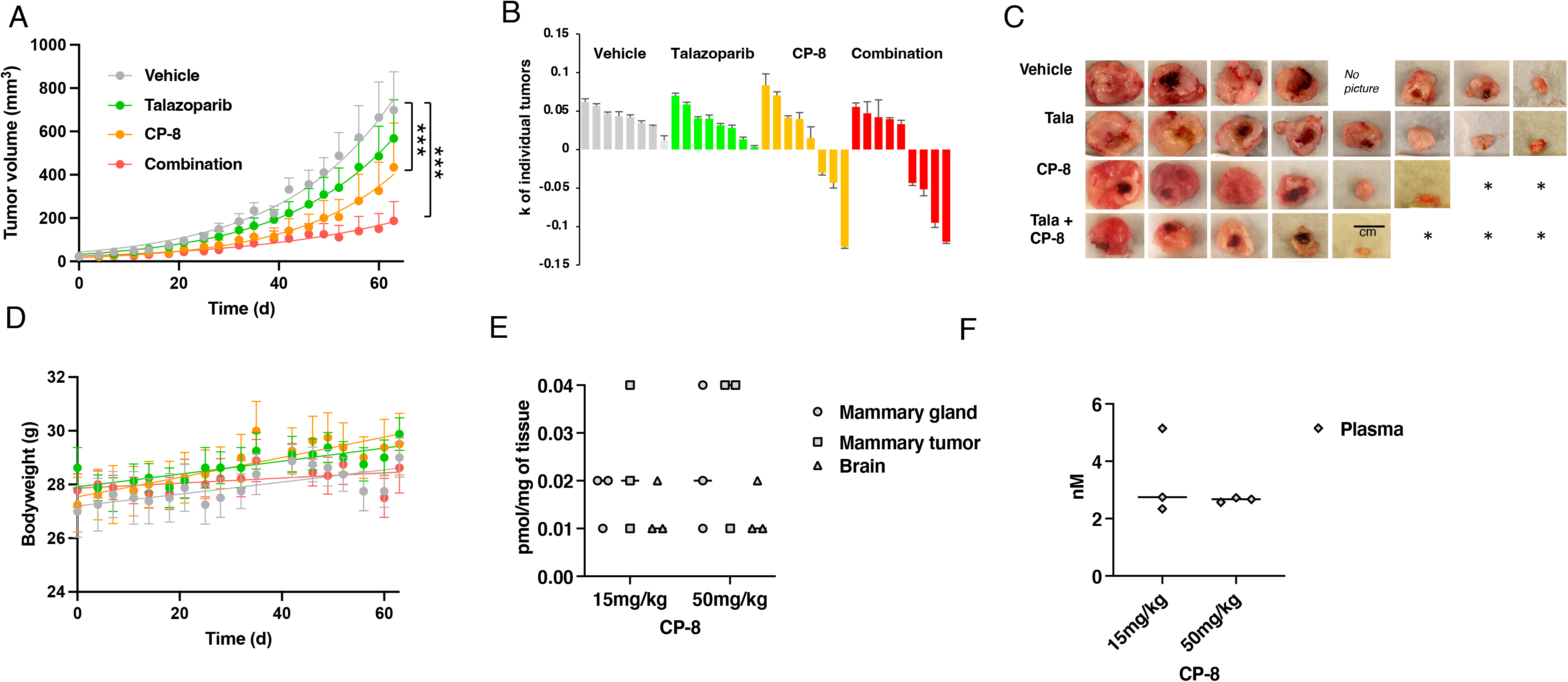
**A.** MM231 xenograft treatment with CP-8 and tala showed tumor reduction with CP-8 alone and in combination with tala. Treatment start: day 0. Vehicle: 7.5% DMSO+61.7% PEG + 30.8% saline. Vehicle n=8, talazoparib n=8, CP-8 n=8 and combination n=9. Statistical analysis was done by Two-way ANOVA, multiple comparisons ***: p<0.0001. **B** k values of the growth curve for each tumor volume over time (day 0 to day 63). **C.** Pictures of tumors collected on day 64. **D.** Weight measurements of mice from (A). **E and F.** Mass spectrometry PK studies in tissues and plasma of 3 SCID mice/group dosed once with 50 mg/kg CP-8.

### CP-8 modulates the oligomeric state of the purified RAD51 protein

The active species of the RAD51 in HR is a nucleoprotein filament formed on ssDNA. Adjacent RAD51 protomers within the filament form the ATPase site, with C319 positioned at this interface near the bound ATP ^10, 29^. The nucleoprotein filament is assembled by adding RAD51 dimer and small oligomers ^30, 31^. We, therefore, expect a similar ATPase active site and protomer-protomer interface organization in RAD51 in the solution. To probe the effect of CP-8 on RAD51 oligomer formation, we used mass photometry, a single-molecule approach that enables the accurate mass measurement of protein molecules in solution ^15^. This analysis allowed us to examine the distributions of RAD51 oligomeric states at concentrations similar to those found in cells.

We compared CP-8 and CP-6 under two reaction conditions that permit (Mg^2+^-ATP) or inhibit (Ca^2^ ^+^-ATP) ATP-hydrolysis by RAD51 nucleoprotein filament ^32^. Under both conditions, RAD51 exists as a mixture of oligomeric species ranging from monomers, dimers, trimers, etc., to much larger oligomers (**Figure 7**). The MP analysis in the presence of inhibitors revealed that in the presence of Mg^2+^-ATP in the reaction buffer, CP-8 promotes RAD51 oligomer formation with distinct peaks at molecular weights representing RAD51 dimer and oligomers containing 10-12 RAD51 monomers, while under Ca^2+^ conditions, CP-8 predominantly promotes dimeric RAD51. In contrast, under Mg^2+^ conditions, CP-6 induced peaks that resembled molecular weights of RAD51 dimer, trimer, and higher oligomeric structures of up to 6 RAD51 proteins. Oligomeric structures of higher order were even more prominent in the presence of Ca^2+^ (**Figure 7**).

**Figure 7.**
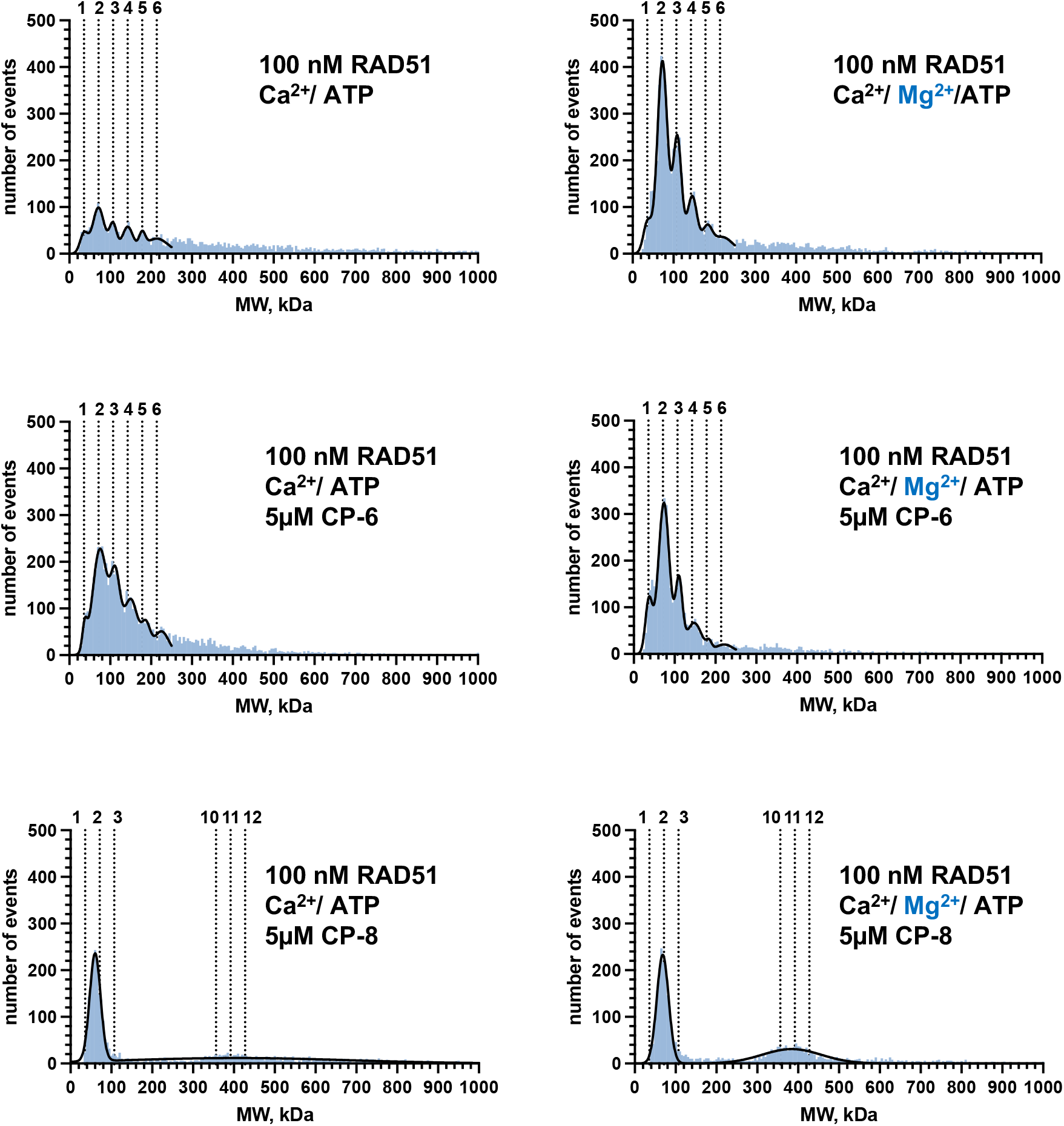
CP-8 changes the distribution of RAD51 oligomers. Mass photometry of RAD51 proteins treated with CP-8 or CP-6 shows the molecular weights of the RAD51 oligomers in solution binned into 5 kDa bins and plotted as histograms (blue). The binned data were then fitted to a sum of Gaussians (black line). Dotted lines indicate the molecular weights of individual oligomeric species. n=2

### RAD51 Cys319 is a target for nitro-alkylation by CP-8 and effectively kills TNBC patient-derived organoids (PDOs)

We have recently shown that Cys319 in RAD51 is a redox-sensitive residue critical for RAD51 function in cells ^10^. Replacing RAD51 Cys319 with Ser using CRISPR/Cas9 in MDA-MB-231 cells significantly reduces cell proliferation (**Figure 8A**) ^10^. Nitroalkenes react with the cysteine thiolate. Thus, unsurprisingly, exposing Cys319Ser cells to increasing concentrations of CP-8 significantly increased CP-8 EC50 values by 2.5 fold (**Figure 8B**). Combined with our data demonstrating that cysteine oxidation is also required for the radio-sensitizing effects of CP-8 (**Figure 4**), this supports the hypothesis that RAD51 Cys319 alkylation is required for the anticancer effects of nitroalkenes. Lastly, to test CP-8 efficacy as monotherapy, we treated TNBC PDOs for 3 days daily with 5 μM or 10 μM CP-8 and assessed. As shown in **Figure 8C**, 10 μM CP-8 effectively decreased PDO viability by 67.1% and 64.3%, respectively.

**Figure 8.**
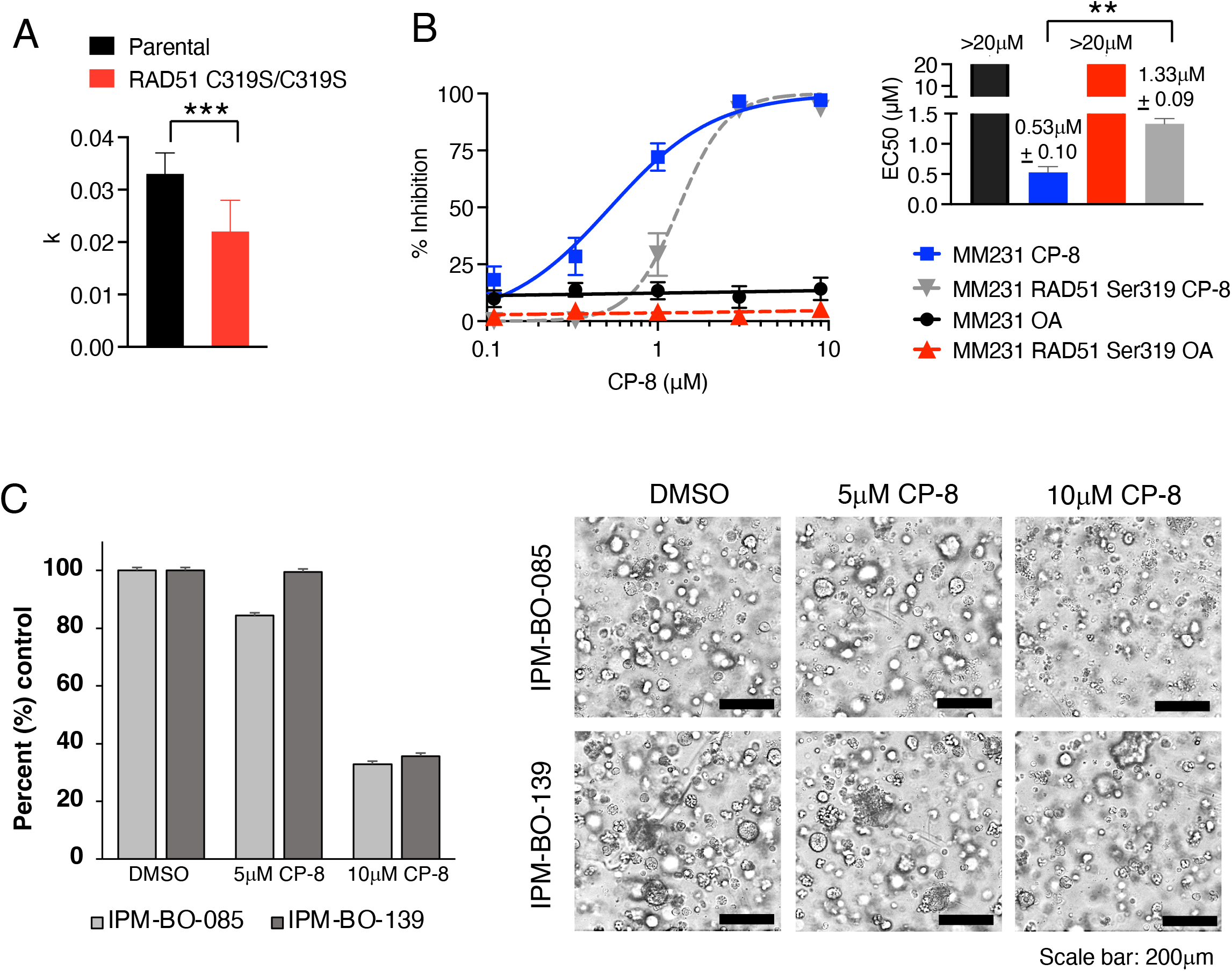
A. Replacing Cys319 with Ser (CRISPR)^10^ increases the EC50 of CP-8. Cells were counted every 4 h for 6 days. K values of the logistic growth curve are plotted and analyzed by Two-way ANOVA. **B.** increases the EC50 of CP-8 in MM231 cells in a 3 d CellTiterGlo assay. OA = oleic acid, a non-nitrated nitroalkene control. **C**. TNBC organoids were treated daily with 5 μM and 10 μM CP-8 for 3 days. Viability was assessed with CellTiter Glo3D (relative luminescence), and data were normalized to vehicle (DMSO) treated cells (left). Bright-field microscopy images showing representative phenotype of treated PDOs (right). Average ± SEM n=3.

## DISCUSSION

Out of all breast cancers (BC), ∼20% are triple-negative breast cancers (TNBC) devoid of the three receptors that define breast cancer treatment strategies such as estrogen receptor (ER), progesterone receptor (PR), and ERBB2 (also known as HER2). TNBC is a very heterogenous cancer characterized by complex genomes driven by genomic instability, copy number alterations, and chromosomal rearrangements 19-23. Unlike other BC, TNBC disproportionally affects younger women and those of African origins and recurs early, within five years after diagnosis, in ∼ 40% of the patients ^33, 34^. Standard care for TNBC patients includes surgery, radiation treatment, and chemotherapy. ^35–39^

Consistently, TNBC treatment has been guided by defects in double-strand DNA repair because homologous recombination (HR)-deficiency induced by germline or somatic loss of the *BRCA1/2* genes, have long been known to sensitize TNBC to DNA damaging agents, including PARP inhibitors (PARPi)^35–37^ The continued relevance of this insight is evidenced by the recent FDA approval of PARPi (olaparib and talazoparib) in metastatic TNBC patients carrying germline mutations in *BRCA1/2* genes ^40^. The incompatibility of simultaneous *BRCA1/2* deficiency and PARPi with cancer survival is the paradigmatic example of synthetic lethality and has motivated research into identifying other synthetic lethal relationships to exploit therapeutically^40–43^. Indeed, the concept of “BRCAness” has emerged to denote any cancer with HR deficiency (HRD) that is sensitive to PARPi or other DNA damaging agents, even in the absence of *BRCA1/2* deficiency^39^. The goal is that by identifying genetic mutations that confer “BRCAness” or inducing it pharmacologically, the landscape of PARPi-sensitive disease can be expanded to include other TNBC patients with limited treatment options^39,41–43^. Recently, olaparib was FDA-approved as *an adjuvant* treatment for patients with germline BRCA-mutated HER-2 negative high-risk early breast cancer ^41^. While exiting, this new treatment option emphasizes research into problems stifling optimal clinical usage of PARP inhibitors (PARPi) because most PARPi-treated *gmBRCA1/2* patients develop resistance due to restoration of HR-mediated DNA DSB repair ^42–44^ and PARPi treatment-associated toxicities that require dose adaptations or drug discontinuation ^45–49^. In addition to these limitations, only 15-20% of TNBC patients are positive for *gmBRCA1/2* and currently receiving PARPi therapy, preventing 80% of TNBC access to PARP inhibitor therapy. Besides surgery, patients with TNBC wildtype BRCA1/2 are usually treated with DNA-damaging therapies such as cisplatin and IR or the microtubule inhibitor paclitaxel ^50–52^.

We recently reported that CP-6 (OA-NO_2_) sensitizes various BRCA-proficient TNBC cell lines to DNA-damaging agents, including doxorubicin, cisplatin, olaparib, and IR ^7^. We screened a library of compounds structurally related to CP-6 to improve nitroalkene efficacy. We demonstrate that the lead candidate CP-8, one carbon longer with the nitroalkene group, shifted two-carbon closer to the carboxylic acid than CP-6. Our studies revealed that compared to CP-6 and the other lead candidate, CP-23, CP-8 shows higher efficacy in targeting HR-proficient human and murine TNBC cells as a single agent and synergistically in combination with IR and PARPis, with the latter currently being FDA-approved only for BRCA-mutant TNBC. Our in vitro and in vivo results strongly suggest that CP-8 cotreatment will allow the inclusion of the ∼80% of TNBC patients expressing wildtype BRCA1/2 genes into PARP-inhibitor treatment.

Besides CP-6 showing action as a RAD51 inhibitor through specifically targeting RAD51 Cys319 ^7^, early human clinical studies of CP-6 (named CXA-10) deemed it as a safe oral renoprotective agent for treating deoxycorticosterone acetate-salt nephropathy with favorable pharmacokinetics ^53, 54^. Nitrated lipids are present in healthy human plasma and urine concentrations of 2 and 20 nM, respectively ^55–57^. These levels can be modulated by diet, reaching plasma levels of up to 20 nM ^58^, paralleling levels achieved after pharmacological interventions using OA-NO_2_^54^. Generated by digestive- and inflammatory-induced nitration reactions, endogenous and therapeutic concentrations of nitroalkenes alkylate only a limited number of hyper-reactive proteins at Cys residues residing in motifs conferring a high degree of Cys nucleophilicity and steric amenability to Michael addition^59^. Notably, the cysteine thiol reactome differs between normal and cancer cells ^60–65^. Therefore, based on our results, we assume that RAD51 Cys319’s reactivity with CP-8 is enhanced in the cancer cells compared to normal cells and tissues examined here. However, future studies need to confirm this in greater detail.

Greater than 10% of FDA-approved small molecule drugs are Michael acceptors (e.g., Omeprazole, Tecfidera, Clopidogrel) or are metabolized to electrophilic products and are considered safe^66^. Nevertheless, there is a reluctance to pursue nitroalkenes as drug candidates because of their presumed unselectivity, which may cause toxicity. Our data convincingly show that CP-8 has specific anti-tumor activity *in vitro* and *in vivo* in murine and human cells that is enhanced by DNA-damaging therapies. Furthermore, while other CP-8 targets conveying the anti-tumor response most likely exist, HR inhibition through RAD51 Cys319 oxidation, based on data presented here, is a significant component of this response. Additional targets need to be identified in future studies.

Like other fatty acids, nitro fatty acid metabolism produces saturation, α-, β- and ω-oxidation products, form co-enzyme A conjugates, and are incorporated into triglycerides and phospholipids ^67, 68^. CP-6 can be shortened through β-oxidation to dinor (minus two carbons), tetranor (minus four carbons), or hexanor (minus six carbons) forms ^69, 70^. Thus, given its metabolite-like feature, we initially expected that CP-23, an even shorter metabolite (C8), would reach high accumulation in tissues and plasma. However, while in vitro data encouraged testing of CP-23 in vivo, PK studies revealed that CP-23, a dimethyl ester 8-carbon long nitroalkene, was predominantly found in mammary tissue and plasma in a partially hydrolyzed forms such as mono-methyl/mono-carboxy or di-carboxy metabolite which may explain its weak performance in the TNBC mouse study. This was surprising given similar demethylation processes underlying the metabolism of the diester and prodrug di-methyl-fumarate (DMF) to mono-methyl fumarate (MMF), which does not affect efficacy. Compared to DMF, MMF is much less susceptible to esterase hydrolysis and spontaneous hydrolysis, preventing further gastrointestinal degradation and enabling absorption into the pre-systemic circulatory system ^71,72^.

CP-8, on the other hand, yielded robust levels in plasma and mammary tissues (benign or cancerous) and the brain, documenting that CP-8 successfully passes the brain-blood barrier. This is important as PARPi are increasingly considered for treating brain metastasis in TNBC patients ^73^. As expected, we found common long-chain fatty acid metabolites of CP-8, such as dinors and tetranors, and reduced forms of CP-8 and related dinors and tetranors, both esterified and free. Further detailed PK and PD studies need to examine the efficacy of the non-reduced esterified and shortened CP-8 isoforms in inhibiting TNBC tumor growth. CP-8 displayed dose-dependent linear increases in plasma and tissue levels after oral 15mg and 50mg/kg doses, without detrimental effects on overall weight gains and indicating any toxicity. This suggests a similar favorable safety and toxicity profile CP-8 and CP-6, making nitro fatty acids attractive drug candidates compared to irreversible inhibitors ^74^, with less toxicity due to the reversibility of their reactions with protein cysteines compared to non-reversible inhibitors ^75,76^.

ATP binding and hydrolysis by the RAD51 nucleoprotein filament are sensitive to solution conditions. In general, Ca^2+^ suppresses ATP hydrolysis, thus elongating the RAD51 oligomer on ssDNA, while Mg^2+^ supports ATP hydrolysis, thus promoting ATP hydrolysis and RAD51 ssDNA disassembly resulting in shorter RAD51 oligomeric structures ^32, 77, 78^. This suggests that both cations may affect RAD51 oligomerization on ssDNA differently. Corroborating this, flow linear dichroism revealed that in the HsRad51/single-stranded DNA filament, the primary intermediate of the strand exchange reaction, ATP/Ca^2+^ induces an ordered conformation of DNA, with a preferentially perpendicular orientation of nucleobases relative to the filament axis, while the presence of ATP/Mg^2+^, ADP/Mg^2+^ or ADP/Ca^2+^ does not ^79^. In the absence of ssDNA or dsDNA, purified human RAD51 displays very little ATPase activity, and the effect of Ca^2+^ or Mg^2+^ under these conditions is unclear. Our MP data obtained in the absence of ssDNA or dsDNA show that Ca^2+^ and Mg^2+^ affect RAD51 oligomerization differently, which is exuberated by adding CP- 8, after which a very distinct population of RAD51 oligomers appears that contain mainly two RAD51 proteins in the presence of Ca^2+^.

Furthermore, Mg^2+^ promotes another population of oligomers containing 10-12 RAD51 proteins. This strongly suggests that CP-8 locks RAD51 in distinct oligomeric forms, thus probably inhibiting DNA binding dynamics or ATP hydrolysis. Future studies are needed to define the details of these mechanisms. As RAD51 resides not only in the nucleus but also in the cytoplasm from which it translocates into the nucleus after IR-induced DNA damage ^10, 80–82^, it is also possible that CP-8 adduction to RAD51 negatively affects RAD51 translocation, which would also support the phenotypes described here demonstrating reduced RAD51 foci formation after IR. Future studies will need to address this in more detail.

In conclusion, the results here encourage using CP-8 as a single agent and as a non-toxic cotreatment with DNA-damaging anticancer therapy. This is important as DNA-damaging therapies such as ionizing radiation, platinum salts, anthracyclines, and PARPi cause toxicities in patients that often require dose adaptations or drug discontinuation ^45–49^. Therefore, we view nitroalkenes like CP-8 as a new drug class of high clinical relevance in anticancer therapy.

## MATERIAL AND METHODS

### Chemical synthesis

Two routes synthesized compounds. CP-8 was synthesized by a one-pot nitro selection reaction of 7-octadecenoic acid (NuCheck Prep), which directly converts an alkene to a non-regiospecific nitroalkene (combination of 7- and 8-nitrononadec-7-enoic acids). Synthesis, workup, and purification were described in published reports for CP-6 ^83^. Dimethyl CP-23 was synthesized by a stepwise condensation method of nitroaldol addition, hydroxyl activation, and elimination to form a regiospecific nitroalkene ^83^. The starting materials methyl 4-nitrobutyrate and methyl 4- oxobutanoate were obtained commercially (Sigma-Aldrich). Compounds CP-24 to CP-30 were formed by transesterification of dimethyl CP-23, refluxing it in the appropriate alcohol with a catalytic amount of PPTS until we observed a complete reaction by HPLC.

### Cell culture

As indicated above, MDA-MB-231 and MDA-MB-468 cells (American Type Culture Collection) were cultured in DMEM growth media. MDA-MB-231 C319S cells mutated via CRISPR/Cas-9 were previously described^10^. MDA-MB-231 and 4T1 cells (American Type Culture Collection) were cultured at 37 °C with 5% CO_2_ in Dulbecco’s modified Eagle’s medium (Gibco) supplemented with 5% FBS (HyClone), 100 units/ml penicillin, 100 mg/ml streptomycin (Gibco), nonessential amino acids (Gibco), and 2 mm L-glutamine (Gibco). MCF10A cells were cultured in DMEM/F-12 (Gibco), 2% horse serum (Gibco), 10 µg/ml insulin (Invitrogen), 0.5 µg/ml hydrocortisone (Sigma-Aldrich), 100 ng/ml cholera toxin (Sigma-Aldrich), 20 ng/ml EGF (Preprotech), 100 units/ml penicillin, 100 mg/ml streptomycin (Gibco). hTERT HME1 cells were cultured in MEGM BulletKit medium (Lonza) and supplemented with 5% FBS for dose-response assays.

### Dose-response assays

Oleic acid (OA) and nitroalkenes were diluted in DMSO (0.05%) and added to cells after solvation in the assay medium. Relative cell numbers were compared by measuring the luminescence signal generated by ATP using the CellTiter-Glo (Promega) assay. Cells were plated in 96-well cell culture-treated plates at 500 cells/well and treated with control OA or nitroalkene analogs (0-6 µM) on days 0, 2, and 6 the following day. For combination drug studies, DMSO or PARP inhibitors talazoparib (0-3.1 µM), olaparib (0-50 µM), or niraparib (0-50 µM) and control OA or nitroalkenes (0-6 µM) for 7 days. Synergy was determined using SynergyFinder^84^.

### Irradiation

Experiments were conducted on cells dosed with 0–10 Gy utilizing a Gammacell 40 Exactor L- Irradiator (Best Medical) with a dose rate of 69 R/min.

### Clonogenic assay

MCF10A (2,000 cells/well) or MM231 cells (1,000 cells/well) were plated on 35mm tissue culture-treated plates and irradiated with 0 to 8 Gy. Following irradiation, cells were treated with 4 µM CP-8 or CP-23 for 24 h. The media was exchanged with culture media, and the plates were incubated for 7 days, with fresh media added every 2 days. After 7 days, colonies were formalin-fixed, stained with crystal violet, and quantified with the ImageJ ColonyArea plugin^85^.

### Cell cycle analysis

MM231 cells were treated with vehicle or 4 µM CP-8 or CP-23 for 24 h and then processed for cell cycle analysis as previously described^10^.

### Homologous Recombination assay

U2OS DR-GFP cells were transfected with I-SceI plasmid to induce DSBs. Six hours later, transfection complexes were removed, and cells were treated with vehicle or 1 μM CP-8 or 1 μM CP-23 with drug replenishment after 24 hours. Forty-eight hours after the transfection drug was removed, HR activity was determined as described previously^7^.

### RAD51 foci determination

MDA-MB-231 cells (150,000 cells/coverslip) were incubated on glass coverslips for 16 h in 6- well plates overnight with culture media. Culture plates were then dosed with 5 Gy ionizing radiation and treated with vehicle, 1 µM CP-8 or 1 µM CP-23 30-45 min after irradiation and processed 6 h later. RAD51 foci formation was detected by immunofluorescent confocal microscopy as previously described^10^.

### Animals

Animals used for this study were approved by and conducted according to the University of Pittsburgh Institutional Animal Care and Use Committee guidelines. MDA-MB-231 cells (1 × 10^6^) were injected into the mammary fat pad (left fourth gland) of 6-week-old female nude mice in a volume of 20 μl of sterile saline as previously described^7^. Eight animals were tested per group. For experimental animals tested with CP-8: When tumors reached an average volume of 30 mm^3^, mice were randomized into groups and administered vehicle (7.5% DMSO+61.7% PEG+30.8% saline), 0.1 mg/kg talazoparib, 15 mg/kg CP-8 or 0.1 mg/kg talazoparib in combination with 15 mg/kg CP-8 every day by gavage (200 μl) for 8 weeks. For experimental animals tested with CP-23: When tumors reached an average volume of 100 mm^3^, mice were randomized into groups and administered vehicle (7.5% DMSO+61.7% PEG+30.8% saline), 50 mg/kg olaparib, 15 mg/kg CP-23, 50 mg/kg CP-23 or combinations of 50 mg/kg olaparib and 15 mg/kg CP-23 or 50 mg/kg olaparib and 50 mg/kg CP-23 every day by gavage (200 μl) for 4 weeks.

### Tissue and plasma processing for CP-8 quantification

Tissue samples were pulverized in dry ice using a tissue pulverizer (Cellcrusher, OR). The resulting powder (between 9-20 mg) was resuspended in 100 μl water and spiked with 100 fmoles of ^15^NO_2_-d_4_-OA as the internal standard, and free and total CP-8 were quantified using HPLC-MSMS. For esterified CP-8, acid hydrolysis was performed by incubating samples with 800 µl of acetonitrile and 100 µl of fuming HCl at 90°C for 1 h. After hydrolysis, 500 µl of an aqueous saturated NaCl solution was added, followed by 1 ml of ethyl acetate. Samples were vortexed and centrifuged at 1,000*g* for 5 min at 4°C. The organic phase was dried under a stream of N2 and reconstituted in acetonitrile/ethyl acetate (70/30, v/v) for quantification by HPLC-MS/MS.

### Chromatographic analysis and quantification of CP-8

Quantification CP-8 and its metabolites were performed in multiple reaction monitoring (MRM) mode following the charged loss of NO2- (m/z 46) or 15NO2- (m/z 47) upon CID using a QTrap 6500+ triple quadrupole mass spectrometer (Sciex, San Jose, CA) equipped with an electrospray ionization source. CP-8 and its metabolites in plasma and tissues were chromatographically resolved by HPLC-ESI-MS/MS using gradient solvent systems consisting of water containing 0.1% acetic acid (solvent A) and acetonitrile containing 0.1% acetic acid (solvent B) using a reverse phase HPLC column (2 × 100 mm 5 µm Luna C18(2) column; Phenomenex) at a 0.70 ml/min flow rate. Samples were loaded onto the column at 40% B, maintained for 0.3 min, and eluted with a linear increase in solvent B from 40-100% of B over 7 min.

CP-8 levels were measured in mice with tumors by dosing mice with CP-8 by gavage, then 2 h later, processing plasma and tissue for quantification. The mice were exsanguinated under isoflurane anesthesia, and the mammary gland, brain, and xenograft tumors were snap-frozen in liquid nitrogen. Tissues were pulverized, and homogenates (250 mg/ml) were resuspended in phosphate buffer (pH 7.4) and incubated with 20 mM mercury (II) chloride (HgCl2) for 30 min at 37°C before lipid extraction.

### Tissue and plasma processing for CP-23 quantification

Mice (3 mice per group) were dosed with 100 mg/kg CP-23 by gavage. After 2 or 4 h, mice were exsanguinated under isoflurane anesthesia, and mammary tissue was harvested and snap-frozen in liquid nitrogen. As CP-23 does not efficiently ionize in ESI, a method based on its derivatization with tiopronin was used ^86^. 50 µl of plasma were mixed with N-(2- Mercaptopropionyl)glycine (tiopronin, Sigma-Aldrich) solution to reach a final concentration of 100 mM and incubated for 30 minutes at 37°C. ^15^NO_2_-d_4_-OA (100 moles) was added as an internal standard. CP-23 derivatives were extracted using 200 µl of cold acetonitrile. Frozen tissue samples were pulverized using a cell crusher. 50 µl of phosphate buffer pH 7.4 (100 mM) was added to 30 mg of powder and spiked with 150 fmoles of ^15^NO_2_-d_4_-OA as internal standard. Tiopronin was added at a final concentration of 100 mM, incubated for 30 minutes at 37°C, and extracted using 200 µl of cold acetonitrile. HPLC-ESI-MS/MS quantified Cp 23 derivatives.

### Chromatographic analysis and quantification of CP-23

Species were chromatographically resolved using an HPLC column (2 × 100 mm 5 μm Luna C18(2) column; Phenomenex) at a flow rate of 0.70 mL/min and a gradient solvent system consisting of Solvent A water containing 0.1% acetic acid, and solvent B acetonitrile containing 0.1% acetic acid. Samples were loaded onto the column at 10% B, maintained for 0.3 min, and eluted with a linear increase in solvent B from 10 to 100% over 4 min. Mass spectrometry was performed in negative ion mode using a QTrap 6500+ triple quadrupole mass spectrometer (Sciex, San Jose, CA) equipped with an electrospray ionization source. Analyte quantification was performed in multiple reaction monitoring (MRM) mode following the charged loss of tiopronin (m/z 162) in negative ion mode upon CID. The following MRMs were used for CP-23 (407/162), mono methyl CP-23 (393/162), dicarboxylate CP-23 (379/162), and ^15^NO_2_-d_4_-OA (494/162)

### Cell cycle DNA content

Propidium iodide (PI) stained DNA content was measured to indicate the cell cycle phase in PRDX1-proficient and deficient MDA-MB-231 cells. 1x10^6^ unsynchronized cells were harvested by trypsinization, and following the inactivation of trypsin, cells were pelleted and washed with ice-cold 1xPBS (Gibco). Cells were fixed in 70% ethanol by adding ethanol dropwise with swirling and incubated for 20 min at 4 °C. Cells were then centrifuged at 2000 rpm, and the supernatant was removed. Pellets were washed with ice-cold PBS and then treated with 500 μl RNase A (50 ng RNAase/μl) for 15 min at 37 °C. Next, propidium iodide (Sigma) (50 ng PI/μl) was added and incubated for 30 min at 4 °C in a dark chamber. Samples were later analyzed at the Flow Cytometry Core at MWRI utilizing a BD LSRII (BD).

### Mass photometry

Recombinant human RAD51 protein was purified as previously described ^87^. Mass photometry (MP) experiments were performed using the Refeyn TwoMP mass photometry instrument (Refeyn Ltd. Oxford, UK) in buffer containing 20 mM Tris pH 7.5, 150 mM KCl, 10 mM MgCl_2_, 1 mM ATP, and in the presence or absence of 5 mM CaCl_2_. Cover slides were cleaned by sequential washing with miliQ water and 100% isopropanol twice, then with miliQ water, and subsequently dried under an air stream. Silicon buffer gaskets were rinsed sequentially with miliQ water, isopropanol, and miliQ water and dried at room temperature. Dried silicon gaskets were applied to the glass slide with mild pressure and mounted on the mass photometer. Two protein oligomer solutions, β-amylase (56, 112, and 224 kDa) and Thyroglobulin (670 kDa), were used for molecular weight calibrations. In each experiment, 400nM of RAD51 was incubated with 20µM of respective inhibitor for 45 or 90 minutes at room temperature. The protein-inhibitor solution was then diluted 4 times into the buffer-filled gasket yielding a final concentration of 100 nM RAD51 and 5 µM inhibitor. Individual molecular weights collected from 3000 frames (59.9 seconds) were binned in 5kDa bins, plotted as frequency histograms, and fitted to multiple Gaussians in GraphPad Prism.

### Organoid culture and drug treatment

TNBC organoids (IPM-BO-085 and 139) were established by the Institute for Precision Medicine (Pittsburgh, PA) and maintained in media as detailed in ^88^. For dose-response assays, organoids were dissociated to single-cell suspension with Trypsin and plated in 190μL of organoid media at 20000 cells/well in 96-well round bottom plates. Treatments were added 3 days post seeding in an additional 100μL of organoid media containing DMSO (0.05%) or DMSO plus CP-8. Organoid images were obtained with an Olympus IX83 microscope at 4x magnification. Media was collected on day 3 to quantify cell viability by CellTiter-Glo 3D.

### Statistical analysis

Unless otherwise noted, data represent the mean ± SEM from three or more independent experiments. Non-linear curves were generated in GraphPad Prism 7.0 (GraphPad Software, La Jolla, CA, USA) for statistical analysis. EC50 values and standard error were calculated utilizing a non-linear dose-response variable slope model. One-way ANOVA tested significance for multiple groups with Tukey posttest for multiple comparisons between groups or by t-test when groups were less than three.

## Acknowledgments

This work was supported by National Institutes of Health P30CA047904 (Hillman Cancer Center, C.), NIH R56 CA233817 (CAN, BAF), Congressionally Directed Medical Research Programs Breast Cancer Research Program BC180467 (CAN, BAF), NIH R01 GM125944 (FJS), NIH R35GM131704 (MS). The authors would like to thank the Institute for Precision Medicine (IPM), a partnership of the University of Pittsburgh and UPMC, for providing the breast cancer patient derived organoids used in these studies.

