## Supplemental Figures for "Small molecule nitroalkenes inhibit RAD51-mediated homologous recombination and amplify triple-negative breast cancer cell killing by DNA-directed therapies"

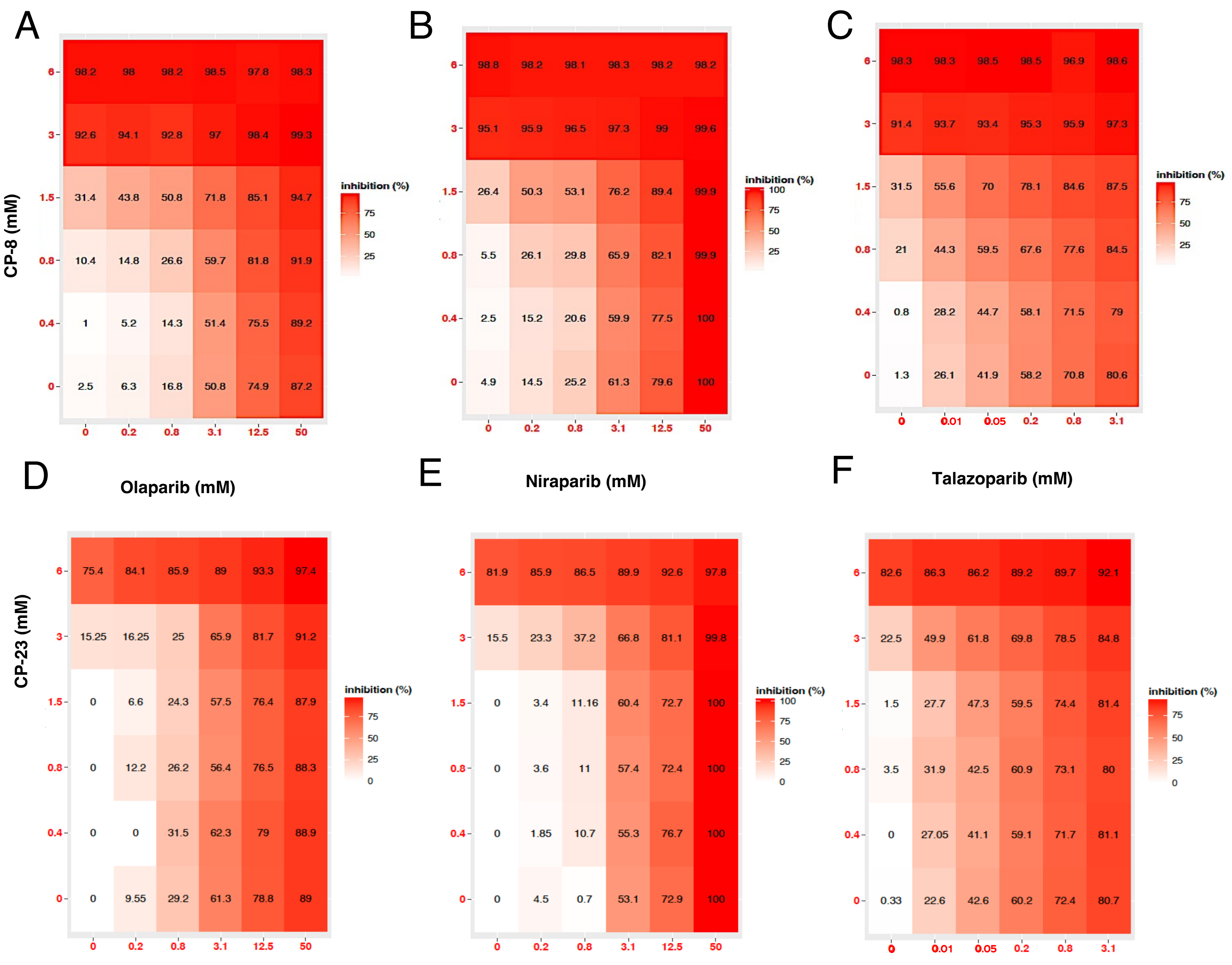


Supplemental Figure 1: A-F. Growth inhibition

### Supplemental Figure 2

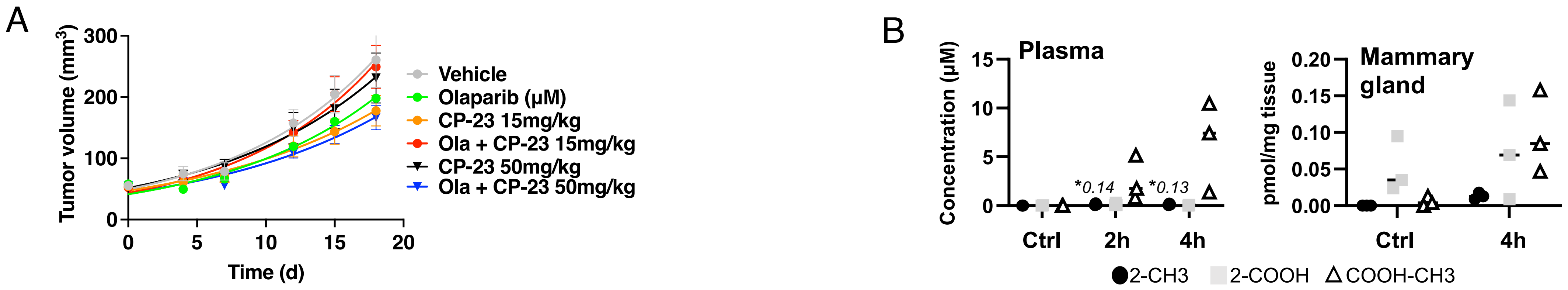


**Supplemental Fig. 2. A.** MM231 xenograft treatment with CP-23 by oral gavage (15 or 50mg/kg and olaparib showed no significant tumor reduction with CP-23 alone or in combination with olaparib. Treatment start: day 0 when tumors were ~60mm$^3$. Vehicle: 7.5% DMSO+61.7% PEG + 30.8% saline. **B** Mass spectrometry PK studies in tissues and plasma of 3 SCID mice/group dosed once with 100mg/kg CP-23. 2-CH3: dimethylated, 2-COOH: di-carboxylated, COOH-CH3: mono-methylated mono-carboxylated.

Supplemental Figure 3

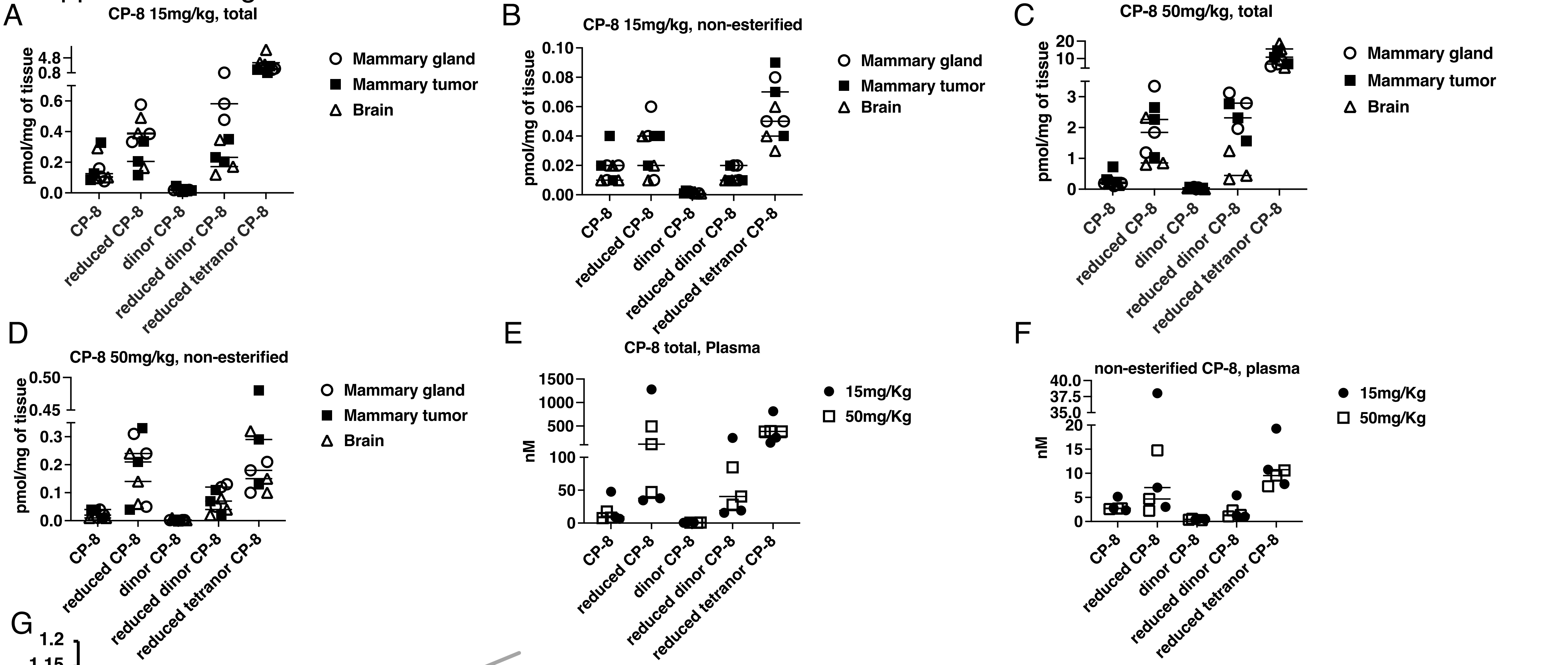


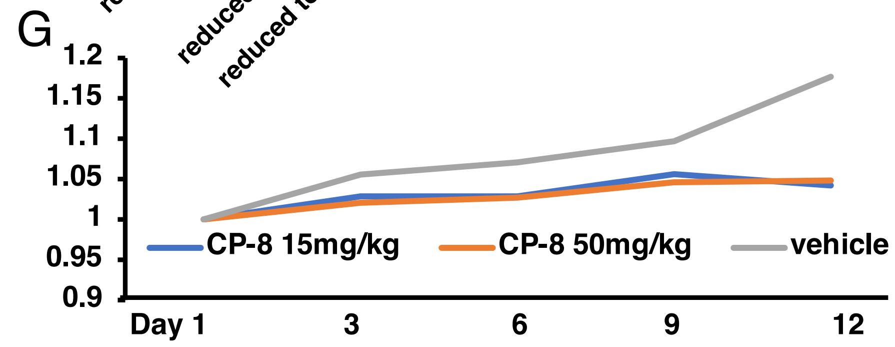


**Supplemental Fig. 2. A –F.** Mass spectrometry PK studies in tissues and plasma of 3 SCID mice/group dosed once with 15 or 50mg/kg CP-8. As expected, common long-chain fatty acid metabolites of CP-8, such as dinors (compounds with 1 round of beta-oxidation) and tetranors (compounds with 2 rounds of beta-oxidation), as well as reduced forms of CP-8 and related dinors and tetranors, both esterified and free, were identified in tissues and plasma, increased by ~3 fold when 15mg/kg dose was increased to 50mg/kg **G.** **M**ouse weight of both 15 mg/kg and 50 mg/kg did not change the weight of mice compared to vehicle control for 12 days.
